# “Divergent TLR2 and TLR4 activation by fungal spores and species diversity in dust from waste-sorting plants”

**DOI:** 10.1101/2022.10.10.511688

**Authors:** Anani K Afanou, Sunil Mundra, Eva Lena Fjeld Estensmo, Ine Pedersen, Jens Rasmus Liland, Elke Eriksen, Pål Graff, Tonje Trulssen Hildre, Karl-Christian Nordby, Anne Straumfors

## Abstract

This manuscript presents the results of an exploratory study on the relationships between NFkB (Nuclear Factor Kappa Chain Enhancer of B-cells) response through TLRs activation by dust characterized by fungal spore concentrations and species diversity. Personal total dust samples were collected from Norwegian waste sorting plants and then characterized for fungal spores and fungal species diversity, as well as for other bioaerosol components, including endotoxins and actinobacteria. The ability of the dust to induce an NFkB response by activating Toll-like receptors 2 (TLR2) and 4 (TLR4) in vitro was evaluated as well as the relationship between such responses and quantifiable bioaerosol components. The average concentrations of bioaerosols were 7.23 mg total dust m^−3^, 4.49×10^5^ fungal spores m^−3^, 814 endotoxin units m^−3^, and 0.6×10^5^ actinobacteria m^−3^. The mean diversity measurements were 326; 0.59 and 3.39 for fungal richness, evenness, and Shannon Index, respectively. Overall, fungal OTUs belonging to the Ascomycotina phylum were most abundant (55%), followed by Basidiomycota (33%) and Mucormycota (3%). All samples induced significant NFkB responses through TLR2 and TLR4 activation. While fungal spore levels were positively associated with TLR2 and TLR4 activation, there was a trend that fungal species richness was negatively associated with the activation of these receptors. This observation supports the existence of divergent immunological responses relationship between fungal spore levels and fungal species diversity. Such relationships seem to be described for the first time for dust from waste facilities.

## 2. Introduction

A variety of operations in waste processing can generate significant levels of complex microbial-rich dust with relatively high fractions of fungal particles. Although fungal exposure has been described for such occupational settings as a potent inflammatory stimulus, the relationships with inflammatory markers in workers are often inconsistent (1– 5) except for specific cases with *Aspergillus sp*. (6).

The airborne mycobiome from waste processing plants encompasses wide taxonomic diversity (7–13) as well as structural and metabolic components that can interact synergistically or antagonistically to elicit immune responses as reviewed for farming settings by May et al (14). The use of adequate in vivo or in vitro models is paramount for identifying hazards and predicting the risks associated with such complex exposure (15).

Many in-vitro cell model studies focusing on pro-inflammatory responses and cytotoxicity have provided valuable insights into our understanding of biologically relevant responses initiated by dust from various exposure settings (6, 16–27). However, underlying pathways and mechanisms of these responses remain unexplored. In fact, many triggering and underlying mechanisms begin with the activation of toll-like receptors (TLRs), which are well-described pattern-recognition receptors (PRRs) of the innate immune system (28). These receptors play a key role in the recognition of microbe-associated molecular patterns (MAMPs) and triggering the internalization and the destruction of microbial particles by immune cells. Various fungal cell wall components such as mannoproteins, beta-glucans, membrane proteins and chitin can induce inflammatory responses in the respiratory tract. While such responses are critical for training the immune system against MAMPs, chronic exposure with repetitive cycles of oxidation and inflammation in the airways is associated with chronic airway disease and a decline in lung function over time (18, 29).

Relatively high exposure levels of fungi have been measured in waste management (5, 17, 27, 30–32). Exposure characterization focusing on how fungal spores and species diversity together with other common bioaerosol components activate TLR2 and TLR4 may provide new insights into important biological effects associated with waste treatment exposure. In the present study, we sought to explore how fungal species diversity or fungal spore levels in dust from waste sorting plants influence the activation TLR2 and TLR4. We aimed therefore to 1) quantify the personal exposure to fungi including fungal spores and species diversity as well as endotoxins and actinobacteria, 2) characterize the induction of the NFkB response in-vitro through TLR2 and TLR4 activation, and 3) explore the relationship between exposure parameters, fungal species diversity and the cell-based immunological effects.

## 3. Materials & Methods

### 3.1. Study sites and sample collection

Air samples were collected at four waste recycling plants in Norway. The recycling plants processed a wide range of waste but mainly waste from offices and various industries. Waste processed during our campaign included wooden office furniture and garden materials. Organic waste or materials contaminated by food waste were also delivered to the waste collection point, but not processed in the plants. These were instead loaded onto trucks, after inspection, and transported to other sites for incineration. The following work tasks, which are probably most exposed, were selected for the measurements: (1) waste reception and inspection, (2) material sorting line, (3) material shredding and pressing, (4) machine operator/driving and (5) storage. All employees in reception and control, sorting, shredding, and pressing as well as machine operators took part in the sampling campaign. The workers were organized into two 8-hour shifts with the heaviest workload on the morning shift. Morning full-shift personal air samples from 18 workers were therefore collected in August, September, and October 2017 at all 4 plants. Detailed sample information is summarized in Table S1.

Personal samples were collected simultaneously with three parallel sampling cassettes worn by each participant. Each worker carried a backpack loaded with pumps attached to each set of filter cassettes placed in the worker’s breathing zone. One sampling set was used for quantification of total dust, fungal spores, and actinobacteria levels and a second set was used for metabarcoding. Dust samples were collected by impaction on preconditioned and pre-weighted 25 mm hydrophilic polycarbonate membrane filters with a pore size of 0.8 µm (Merck KGaA, Darmstadt, Germany) mounted in antistatic polypropylene total dust filter cassettes (Pall Laboratories, Port Washington, NY, USA). A third sampling set with 25 mm glass fiber filters (1 m, GF/A, Whatman, UK) mounted in PAS-6 personal air sampler cassettes (van der Wal, J. (1983) was used for the collection of endotoxins. Air pumps (GSA5200, GSA Messgerätebau GmbH, Ratingen, Germany) were operated at an average flow rate of 2.0 (±10%) liters per minute. The total sampling time varied between 3.9 and 7 hours (mean 5.7 hours).

### 3.2. Gravimetric analysis of the dust

The total dust content was determined gravimetrically using a microbalance (Sartorius AG, MC210p, Göttingen, Germany). The sample-loaded filters were acclimatized (temperature of 20±1 C and relative humidity of 40±2%) for 48 hours before being subjected to gravimetric analysis. Two unexposed blank field filters were included, and the average weight was used as blank to adjust the final dust weight. The detection limit corresponded to 0.02 mg/filter.

### 3.3. Dust elution from filter membrane

The filter samples were eluted in 5 ml PBS with 0.1% BSA in 15 ml centrifuge tubes and subjected to 5 min sonication followed by orbital shaking at room temperature for 60 min. The dust suspension was then transferred to a new tube and the process was repeated once with 2 ml PBS with 0.1% BSA for 25 min. The final dust suspension was aliquoted and kept at

-20°C until microscopic analysis and in vitro experiments.

### 3.4. Field emission scanning electron (FESEM) analysis of actinobacteria and fungal spores

For microscopic analysis, 0.5 ml of each aliquoted sample was filtered onto a 25 mm polycarbonate filter (pore size: 0.45 µm, Merck KGaA, Darmstadt, Germany). The filters were then air dried under sterile conditions and mounted onto 25mm diameter aluminum pin stubs (Agar Scientific Ltd., Stansted Essex, UK) using double-sided adhesive carbon discs. The filter samples were coated with 5-6 nm platinum in a Cressington 200HR sputter coater (Cressington Scientific Instruments Ltd., Watford, UK). Coated samples were analyzed using a FESEM as previously described by (33). Briefly, the microscope was operated in secondary electron imaging mode with an acceleration voltage of 15 keV, an extraction voltage of 1.8 kV and a working distance of 10-11 mm. Fungal and actinobacteria spores were identified by morphological features and counted in 100 randomly selected imaged fields at 3000x magnification. The spore counts were given as the number of spores m^−3^ air.

### 3.5. Fungal community characterization

#### 3.5.1. DNA extraction and metabarcoding

DNA was extracted from exposed filters as well as three unexposed controls using the E.Z.N.A Soil DNA Kit (Omega Bio-tek, Norcross, GA, USA). The filters were transferred to disruptor tubes and 800 µL SLX-Mlus buffer was added. The samples were homogenized (2 x 1 min at 30 Hz) by a TissueLyser (Qiagen, Hilden, Germany) and stored at -20 °C until next step. The samples were then thawed at 70 °C, incubated for 10 min and then homogenized again using the same procedure. Thereafter, the samples were chilled on ice before adding 600 µL chloroform. The samples were then vortexed and centrifuged at 13 000 rpm for 5 min at RT. The aqueous phase was transferred to a new tube with an equal volume of XP1 Buffer and vortexed. Samples were transferred to HiBind DNA Mini Column and processed by following the manufacturer’s guidelines. The extracted DNA was eluted in 50 µL elution buffer. The fungal ITS2 region was targeted with the forward primer ITS4 (5′-xCTCCGCTTATTGATATG) (34) and the reverse primer gITS7 (5′-xGTGARTCATCGARTCTTTG) (35), with barcodes x of 6-9 base pairs (bp). The ITS2 region was selected because of less amplification bias due to length differences and the development of less biased primers. The PCR reaction contained 2 µl DNA template and 23 µl master mix containing 14.6 µl Milli-Q water, 2.5 µl 10x Gold buffer, 0.2 µl dNTP’s (25 nM), 1.5 µl reverse and forward primers (10 µM), 2.5 µl MgCl2 (50 mM), 1.0 µl BSA (20 mg/ml) and 0.2 µl AmpliTaq Gold polymerase (5 U/µl, Applied Biosystems, Thermo Fisher Scientific). Negative PCR controls were included.

Amplification was performed by initial denaturation (95 °C for 5 min), followed by 32 cycles of denaturation (95 °C for 30 s), annealing (55 °C for 30 s) and elongation (72 °C for 1 min). A final elongation step was included (72 °C for 10 min). The PCR products were normalized with the SequalPrep Normalization Plate Kit (Invitrogen, Thermo Fisher Scientific, Waltham, MA, USA) and eluted in 20 μL elution buffer. The resulting PCR products were pooled, before concentration and purification by Agencourt AMPure XP magnetic beads (Beckman Coulter, CA, USA). The concentration of the purified pools was determined by Qubit (Invitrogen, Thermo Fisher Scientific, Waltham, MA, USA) before the library was sent to Fasteris SA (Plan-les-Ouates, Switzerland) for barcoding with Illumina adapters, spiking with PhiX and sequencing with 2 x 250 bp paired-end reads with Illumina MiSeq (Illumina, San Diego, CA, USA).

#### 3.5.2. Bioinformatic workflow

Forward and reverse sequences were retrieved from Fasteris SA, which we independently demultiplexed using CUTADAPT v 2.7 (mismatches between barcode tags and sequence primer = 0, minimum sequence length = 100 bp) (36). DADA2 v.12 (37) was used to filter low-quality sequences (maximum expected error=2.5) and to correct errors based on an implemented machine learning model. Then we merged the error-corrected forward and reverse sequences (minimum overlap = 5 bp). Chimeras were filtered out using the Bimera algorithm in DADA2 (default parameters). The resulting amplicon sequence variants were clustered into operational taxonomic units (OTUs; 97% similarity) using VSEARCH (38), before using LULU (39) to correct for possible OTU over-splitting (default parameters). Taxonomy was assigned to the resulting OTU table by BLAST (40) and the UNITE database (41). All negative controls were automatically removed during bioinformatics due to insufficient sequences. The final data set contained 2516 OTUs.

### 3.6. Analysis of endotoxins

Exposed glass-fiber filters were transferred to 15 ml glass tubes and eluted in 5 ml endotoxin-free water containing 0.05% Tween-20 by orbital shaking for 1 hour. The filters were then removed from the tube and the suspension centrifuged at 1000×g for 15 minutes to pellet the particulate fraction. The supernatant underwent a second centrifugation step before being aliquoted without the pellet and stored at -20°C until analysis. For endotoxin quantification, supernatants were diluted 20-50 times before a Limulus amoebocyte (LAL) kinetic-QCL assay was applied according to the manufacturer’s instructions (Lonza Ltd., Basel, CH). The detection limit of the assay was estimated at 0.5 -2.5 EU/filter depending on the dilution rate. Parallel controls consisting of samples spiked with 10 µl endotoxin solution (50 EU ml^−1^) and blank samples were run to assess possible inhibition or enhancement of the sample matrix. The endotoxin concentrations were determined by kinetic measurement of absorbance at 405 nm using a BioTek spectrophotometer (BioTek Instruments Inc., VT, USA) in accordance to five points standard curve with concentrations ranging from 0.005 to 50 EU ml^−1^.

### 3.7. In vitro assay of NFkB response through TLR2 and TLR4 activation

Human embryonic kidney (HEK) 293 reporter cells for TLR2 and TLR4 (Invivogen, France) were exposed to the eluted dust samples to study their potential to induce NFkB responses. The reporter cells specifically expressed TLR-inducible reporter genes encoding for secreted embryonic alkaline phosphatase (SEAP) via the NFkB signaling pathway. SEAP is secreted extracellularly and can be quantified in the supernatant after NFkB induction by TLRs activation. The exposure experiments were performed in a 96-well plate following the procedure described by (42) with some minor modifications. Briefly, we transferred 180 µl per well of 2.8×10^5^ cells ml^−1^ in culture media (DMEM + 10% fetal bovine serum + manufacturers recommended HEK Blue selection antibiotics) and incubated for 3 h at 37 °C (5% CO_2_, high humidity) to allow cells to settle at the bottom and relieve stress. Thereafter, 20 µl of the dust suspension was added in triplicate and the cells were incubated overnight for 22h. The following day, 20 µl of cell supernatant were transferred to new plates and supplemented with freshly made 180 µl Quanti-Blue solution (Invivogen, France). After 180 minutes of incubation, the developed color was measured spectrophotometrically at a wavelength of 649 nm using SpectraMax i3 equipped with SoftMax-Pro 6.3.1 software (Molecular Devices LLC, San Jose CA, USA). We included sample wash buffer (PBS + 0.1% BSA) as a negative control, LTA (lipoteichoic acid; Invivogen, France) as a positive control for TLR2, and ultrapure LPS (lipopolysaccharide; Invivogen, France) as a positive control for TLR4 and Zymosan (Merck KGaA, Darmstadt, Germany) that activate both receptors. Note that a parallel plate with culture media with samples only was included to check absorbance due to dust alone. Final data were presented as arithmetic means of the three replicate measurements adjusted for background from media with samples only.

### 3.8. Data treatment and statistical analysis

Prior to statistical analysis, all concentrations below the limit of detection for fungal spores, actinobacteria and endotoxins were arbitrary replaced by the limit of detection (LOD)/√2 (43). The levels of the exposure components were reported as arithmetic means (AM) with standard deviation and geometric mean (GM) with geometric standard deviation. For comparison between companies, the non-parametric Kruskal-Wallis (K-W) test was used because the data for dust, endotoxins, actinobacteria, and TLR2 were not normally distributed (Shapiro test, *p*≤ 0.05), and because of the small sample size. After a significant K-W-test, post hoc Wilcoxon rank-sum tests (or Mann-Whitney test) with Bonferroni adjustment (*p*≤ 0.0125 for 4 companies) were used. Visualization of TLR activation was performed using bar graphs. In addition, we applied correlation and linear regression analysis to show the relationship between different exposure components and TLR activation. For the multiple comparison tests of exposure agents, a two-sided *p*-value of 0.013 was considered statistically significant. The above statistical analyses were performed using STATA 16 SE (Statacorp LP, College Station, TX, USA).

For fungal diversity, statistical analyses were performed in R v4.0.3 (R Core Development Team 2020). To achieve homogeneity of variances, the absorbance values of the TLR2 and TLR4 activation were transformed to zero skewness and expressed on a 0–1 scale before the analyses, and the fungal community matrix (OTUs table) was arcsine-transformed prior to analyses as used earlier (44). For fungal diversity analyses, the OTUs table was normalized by randomly resampling 10,982 reads per sample. Richness, Shannon Index and Evenness measures were computed using the *Vegan* package and inter-company differences were examined using ANOVA followed by a Tukey’s HSD post-hoc test of the A*gricola*e package and visualized using violins plots. Linear regression analyzes were also performed to examine the relationship between fungal abundance and TLR2/TLR4 activation. We also calculated Pearson’s correlations between fungal genera abundance and TLR4 as well as TLR2 activation absorbances using package *Corplot*. The variation in the abundance of fungal genera different companies was tested using ANOVA (Benjamini-Hochberg FDR correction was applied) followed by a Tukey’s HSD post-hoc test and illustrated by heat plots based on hierarchical clustering. Species accumulation curves for the number of cumulative OTUs for different companies were calculated using Function S*pecaccum* from *Vegan* package (45) with 10000 permutations. The overall taxonomic composition of fungi at phylum level and order level among companies was visualized using stacked bar charts. The Bray–Curtis dissimilarity index was used to generate community distance matrices and further used in all multivariate analyses. To examine the relative importance of different factors (companies) and vectors (TLR2 and TLR4 activation), variables on fungal community structural pattern, multivariate permutational analysis of variance (PERMANOVA) test was used, as implemented in the Adonis function of the package *vegan*. The PERMANOVA analysis was performed using a forward selection practice to improve the final model. We first examined single variable models and then included significant factors in the final model in order of their R^2^ values. In addition, Global Nonmetric Multidimensional Scaling (GNMDS) ordination analysis was used to visualize the effects of studied variables on composition of fungal community using the meta MDS function of the V*egan* package. Vector variables and centroids of the factor variables were fitted to GNMDS plots using the *envfit* function and the *ordiellipse* function was used to plot the 95% confidence intervals (CI) of the different companies.

## 4. Results

### 4.1. Exposure levels of dust, fungal spores, actinobacteria and endotoxins

Workers were exposed to dust concentrations ranging from 0.31 mg m^−3^ to 50.22 mg m^−3^ with a geometric mean and geometric standard deviation GM (GSD) of 1.8 (4.6) mg m^−3^. The arithmetic mean (AM) of dust exposure levels varied significantly between companies (p=0.009) with the highest value measured in Company 1 (AM (median): 13.35 (1.55) mg m^−3^) compared to Company 4 with the lowest concentrations (0.40 (0.36) mg m^−3^) (Table 1). With exception of 3 workers in Company 1, who were exposed to 11.04, 48.55 and 50.22 mg.m^−3^, respectively, the overall dust exposure values were below the Norwegian Occupational Exposure Limit (OEL) of 5 mg m^−3^. The total dust contained relatively high levels of fungal spores (Figure 1A) ranging from below the detection limit to 15.3×10^5^ spores m^−3^. Spore concentrations varied significantly between companies (p=0.009) with the highest measured exposure levels at Company 1 (AM (median): 7.76 (7.45) ×10^5^ spores m^−3^) compared to Company 3 (0.4 (0.04) ×10^5^ spores m^−3^) (Table 1). Airborne actinobacteria were also detected but only in Company 1 (Fig 1B, Table 1) in 4 samples. Relatively high levels of endotoxins were detected in all samples with only 3 samples having values below the recommended health related OEL of 90 EU m^−3^(46). Four workers were exposed to more than 1000 EU m^−3^ while the exposure levels for the remaining workers varied between 90 and 1000 EU m^−3^. The highest values were found in Company 1 (AM (median): 1466 (344) EU m^−3^) versus Company 2 with the lowest levels (123 (126) EU m^−3^), but no statistically significant differences could be found between the companies (Table 1).

**Table 1:** Descriptive statistics of overall data and stratified by companies

| Bioaerosol components | N | AM | SD | MEDIAN | GM | GSD | MIN | MAX |
| --- | --- | --- | --- | --- | --- | --- | --- | --- |
| Dust (mg m<sup>-3</sup>) |  |  |  |  |  |  |  |  |
| Overall | 18 | 7.23 | 15.54 | 1.44 | 1.79 | 4.56 | 0.31 | 50.22 |
| Company 1 | 9 | 13.35 | 20.69 | **1.55<sup>a</sup>** | 3.98 | 5.18 | 0.8 | 50.22 |
| Company 2 | 3 | 2.22 | 0.83 | 1.8 | 2.12 | 1.42 | 1.68 | 3.17 |
| Company 3 | 3 | 0.73 | 0.52 | 0.54 | 0.62 | 2.01 | 0.34 | 1.33 |
| Company 4 | 3 | 0.4 | 0.12 | **0.36<sup>a</sup>** | 0.39 | 1.34 | 0.31 | 0.54 |
| Fungal spores (# m<sup>-3</sup>) |  |  |  |  |  |  |  |  |
| Overall | 18 | 4.9×10<sup>5</sup> | 4.6×10<sup>5</sup> | 3×10<sup>5</sup> | 1.95×10<sup>5</sup> | 7.14 | <LOD | 15.3×10<sup>5</sup> |
| Company 1 | 9 | 7.76×10<sup>5</sup> | 4.52×10<sup>5</sup> | **7.45×10<sup>5b</sup>** | 6.28×10<sup>5</sup> | 2.15 | 1.56×10<sup>5</sup> | 15.3×10<sup>5</sup> |
| Company 2 | 3 | 3.19×10<sup>5</sup> | 3.43×10<sup>5</sup> | 2.67×10<sup>5</sup> | 0.89×10<sup>5</sup> | 15.7 | <LOD | 6.85×10<sup>5</sup> |
| Company 3 | 3 | 0.42×10<sup>5</sup> | 0.67×10<sup>5</sup> | **0.039×10<sup>5b</sup>** | 0.12×10<sup>5</sup> | 7.25 | <LOD | 1.2×10<sup>5</sup> |
| Company 4 | 3 | 2.24×10<sup>5</sup> | 1.09×10<sup>5</sup> | 2.34×10<sup>5</sup> | 2.04×10<sup>5</sup> | 1.75 | 1.1×10<sup>5</sup> | 3.28×10<sup>5</sup> |
| Actinobacteria (# m<sup>-3</sup>) |  |  |  |  |  |  |  |  |
| Overall | 18 | 0.6×10<sup>5</sup> | 1.4×10<sup>5</sup> | 0.04×10<sup>5</sup> | 0.091×10<sup>5</sup> | 5.43 | <LOD | 5.6×10<sup>5</sup> |
| Company 1 | 9 | 1.08×10<sup>5</sup> | 1.8×10<sup>5</sup> | 0.053×10<sup>5</sup> | 0.21×10<sup>5</sup> | 8.31 | <LOD | 5.56×10<sup>5</sup> |
| Company 2 | 3 | 0.041×10<sup>5</sup> | 0.0037×10<sup>5</sup> | 0.039×10<sup>5</sup> | 0.041×10<sup>5</sup> | 1.09 | <LOD | 0.045×10<sup>5</sup> |
| Company 3 | 3 | 0.039×10<sup>5</sup> | 0.01×10<sup>5</sup> | 0.39×10<sup>5</sup> | 0.39×10<sup>5</sup> | 1.03 | <LOD | 0.4×10<sup>5</sup> |
| Company 4 | 3 | 0.37×10<sup>5</sup> | 0.025×10<sup>5</sup> | 0.37×10<sup>5</sup> | 0.37×10<sup>5</sup> | 1.07 | <LOD | 0.4×10<sup>5</sup> |
| Endotoxins (EU m<sup>-3</sup>) |  |  |  |  |  |  |  |  |
| Overall | 18 | 814 | 1441 | 231 | 193 | 8.7 | <LOD | 5321 |
| Company 1 | 9 | 1466 | 1855 | 344 | 283 | 20.7 | 0.35 | 5321 |
| Company 2 | 3 | 123 | 22 | 126 | 121 | 1.2 | 99 | 143 |
| Company 3 | 3 | 180 | 131 | 141 | 150 | 2.11 | 73 | 326 |
| Company 4 | 3 | 183 | 198 | 77 | 124 | 2.84 | 60 | 411 |
| Fungal OTUs richness |  |  |  |  |  |  |  |  |
| Overall | 18 | 326 | 124 | 282 | 306 | 1.4 | 152 | 574 |
| Company 1 | 9 | **276<sup>c</sup>** | 41 | 262 | 273 | 1.2 | 224 | 340 |
| Company 2 | 3 | 303 | 107 | 266 | 291 | 1.4 | 219 | 424 |
| Company 3 | 3 | 270 | 122 | 263 | 251 | 1.62 | 152 | 396 |
| Company 4 | 3 | **557<sup>c</sup>** | 23 | 565 | 556 | 1.04 | 531 | 574 |
| Evenness index |  |  |  |  |  |  |  |  |
| Overall | 18 | 0.59 | 0.12 | 0.60 | 0.57 | 1.31 | 0.23 | 0.76 |
| Company 1 | 9 | 0.56 | 0.06 | 0.56 | 3.11 | 1.12 | 0.42 | 0.64 |
| Company 2 | 3 | 0.61 | 0.04 | 0.60 | 3.47 | 1.14 | 0.58 | 0.66 |
| Company 3 | 3 | 0.51 | 0.25 | 0.62 | 2.54 | 2.01 | 0.23 | 0.70 |
| Company 4 | 3 | 0.72 | 0.03 | 0.71 | 4.55 | 1.04 | 0.70 | 0.76 |
| Shannon diversity index |  |  |  |  |  |  |  |  |
| Overall | 18 | 3.39 | 0.83 | 3.36 | 3.261 | 1.367 | 1.14 | 4.74 |
| Company 1 | 9 | **3.13<sup>d</sup>** | 0.33 | 3.25 | 0.55 | 1.13 | 2.47 | 3.49 |
| Company 2 | 3 | 3.49 | 0.46 | 3.35 | 0.61 | 1.07 | 3.11 | 4.00 |
| Company 3 | 3 | 2.91 | 1.58 | 3.44 | 0.46 | 1.85 | 1.14 | 4.16 |
| Company 4 | 3 | **4.55<sup>d</sup>** | 0.16 | 4.49 | 0.72 | 1.04 | 4.43 | 4.74 |
| TLR2 (absorbance) |  |  |  |  |  |  |  |  |
| Overall | 18 | 0.68 | 0.31 | 0.59 | 0.61 | 1.64 | 0.15 | 1.35 |
| Company 1 | 9 | 0.85 | 0.33 | 0.75 | 0.80 | 1.50 | 0.44 | 1.35 |
| Company 2 | 3 | 0.60 | 0.01 | 0.60 | 0.60 | 1.02 | 0.59 | 0.61 |
| Company 3 | 3 | 0.55 | 0.12 | 0.51 | 0.54 | 1.23 | 0.46 | 0.68 |
| Company 4 | 3 | 0.37 | 0.19 | 0.43 | 0.32 | 1.94 | 0.15 | 0.51 |
| TLR4 (absorbance) |  |  |  |  |  |  |  |  |
| Overall | 18 | 0.56 | 0.21 | 0.52 | 0.52 | 1.52 | 0.17 | 0.94 |
| Company 1 | 9 | 0.70 | 0.18 | 0.70 | 0.68 | 1.30 | 0.45 | 0.94 |
| Company 2 | 3 | 0.48 | 0.07 | 0.44 | 0.47 | 1.16 | 0.42 | 0.56 |
| Company 3 | 3 | 0.47 | 0.12 | 0.43 | 0.47 | 1.27 | 0.39 | 0.61 |
| Company 4 | 3 | 0.33 | 0.16 | 0.32 | 0.30 | 1.71 | 0.17 | 0.49 |

AM: arithmetic mean, SD: standard deviation GM: geometric mean, GSD: geometric standard deviation. LOD: limit of detection. (Kruskal Wallis followed by Wilcoxon rank sum test: a: $p=0.009$ ; b: $p=0.009$ ; c: $p=0.0002$ ; d: $p=0.03$ )

**Figure 1:**
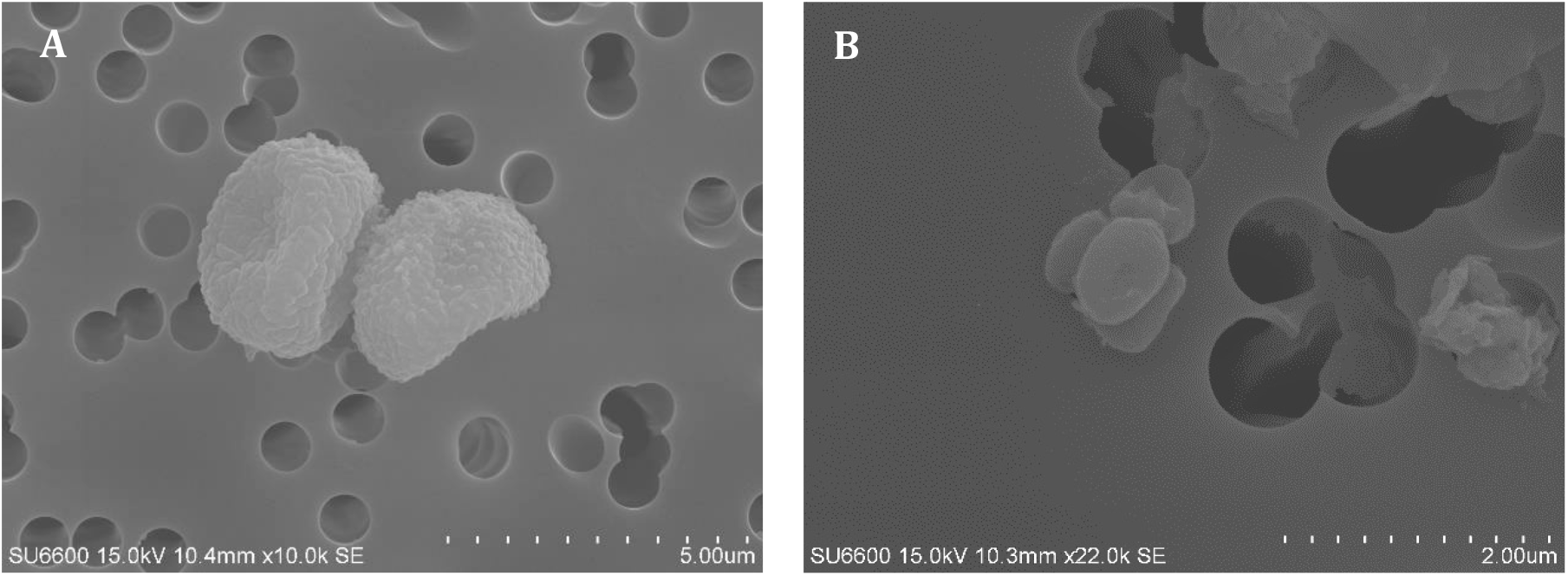
Micrographs of spores from fungi (A) and actinobacteria (B)

### 4.2. TLRs activation by the dust samples

All samples induced significant activation of TLR2 and TLR4 as compared to parental HEK reference cells and negative controls (Figure S1). Three samples with dust levels above 5 mg m^−3^ induced stronger activation of both receptors of positive controls of 10 µg mL^−3^ of LTA and 10 ng mL^−3^ of LPS, respectively. The highest activation values were observed in the dust samples from Company 1 and the lowest activations in the dust samples from Company 4. Activation of TLR2 correlated strongly with the dust levels (r= 0.8, p=0.0001), but weakly with endotoxins (r= 0.42, p=0.08) and fungal spores (r= 0.41, p=0.09). Similarly, activation of TLR4 correlated strongly with dust (r= 0.71, p=0.001) and moderately with endotoxins (r= 0.52, p=0.03) and with fungal spore counts (r= 0.53, p=0.02). This was also supported by linear regression analysis (Table S2). Furthermore, SEAP levels by TLR2 activation were higher than that those by TLR4 activation, with the exception of 2 samples (S1 (Company 1) and S11 (Company 3), indicating the presence of more ligands for TLR2 than TLR4 in the collected samples.

### 4.3. Inhalable mycobiome of the waste management

#### 4.3.1. Fungal diversity patterns

We found significantly higher OTUs richness among workers in Company 4 (AM±SD: 557± 23) versus Company 1 (276±41), 2 (303±107) and 3 (270±122) (Figure S2a). In particular, the sample with the highest richness came from a worker who performed sorting task in Company 4. The lowest diversity was measured among workers in Company 3, and the sample came from a worker who worked with shredders and presses. Notably, the two samples with the highest and lowest richness were collected from companies in the Oslo region in September. Consistent with the richness patterns, the Shannon diversity index and the evenness were higher in samples from workers at Company 4 compared with Company 1, but the differences were not significant (p=0.03 and p= 0.10, respectively (Figures S2b and S2c)). The abundance of the 10 most common OTUs in the data set was highest in companies 1 and 3, and lowest in Company 4 (Figure S2d). However, companies 1 (n=432) and 4 (n=447) had a higher number of OTUs than companies 2 (n=94) and 3 (n=195), and a total of 271 OTUs were common to all companies (Figure S3).

We found no significant linear relationships between fungal diversity measures (i.e. richness, Shannon diversity index and evenness index) and exposure variables (dust, endotoxin, fungal spores and actinobacteria) (data not shown). However, we observed that fungal richness was negatively associated with the activation of TLR4 (R^2^ adj. =0.28, p = 0.014) and weakly with TLR2 (R^2^ adj. = 0.16, p = 0.06) (Figure S4).

#### 4.3.2. Fungal composition by companies and relationship with TLRs activation

Overall, the phylum of Ascomycota (55%) dominated the OTUs reads compared to Basidiomycota (33%) and Mucoromycota1 (2%) (Figure 2; Table S3). By company, Ascomycota represented 69%, 58%, and 59% of reads in Company 1, 2 and 3, respectively, while in Company 4 they represented only 24%. The Basidiomycota dominated the OTU measurements in Company 4 (75%) while they accounted for about 33% in companies 2 and 3 and only 11% in Company 1 (Figure 2a).

**Figure 2:**
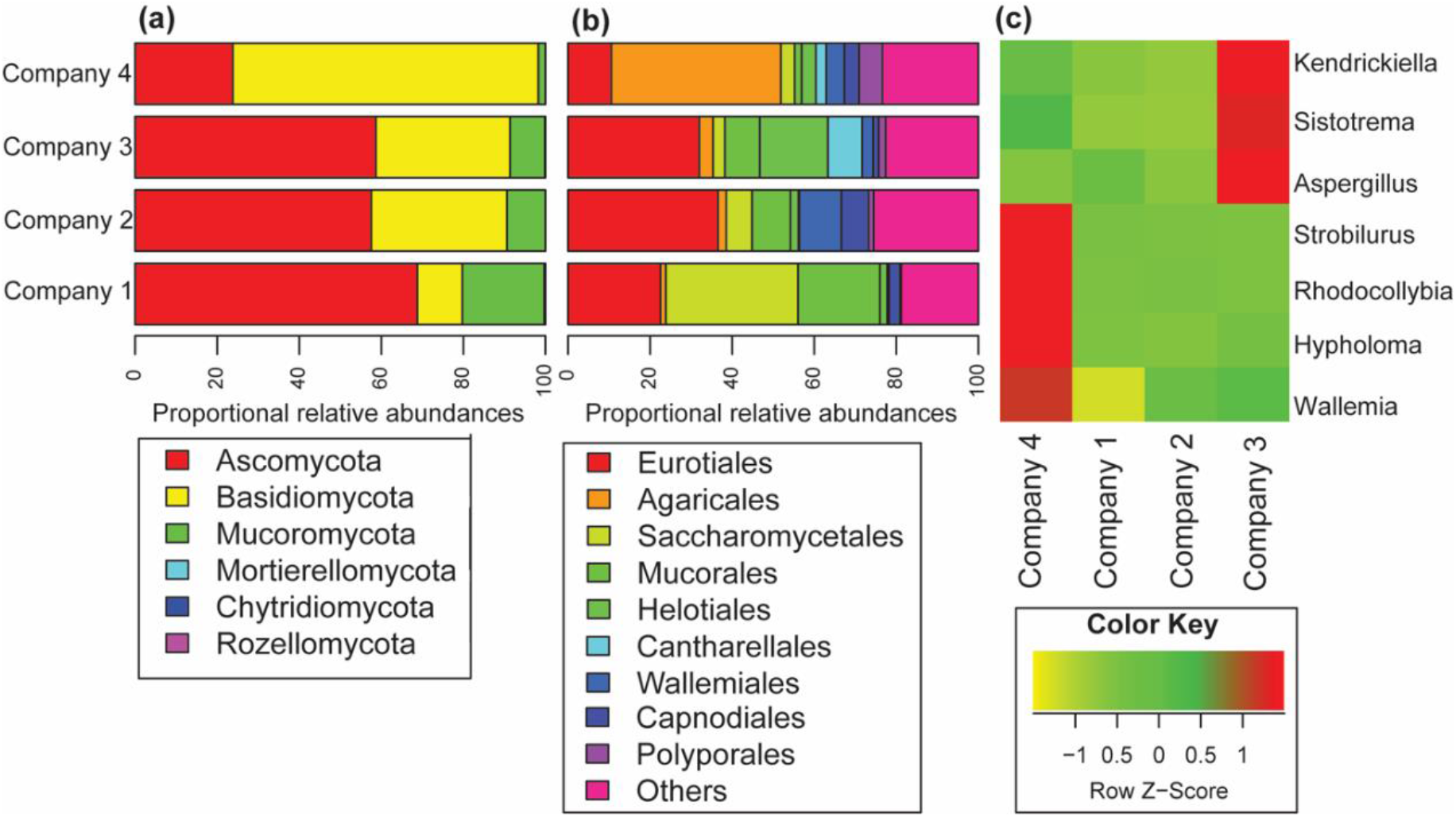
Taxonomic distribution of the fungal communities detected from four recycling plants (company 1 to 4). Proportional relative abundances of airborne fungal compositions at (a) phyla and (b) order level. The data represent average reads per samples. (c) Hierarchical clustering-based heat-plots for proportional abundances of different genera that are significantly varying in their abundance among the four companies.

The phylum Mucoromycota (Zygomycota) accounted for 20%, 9%, 9% and 2% in company 1, 2, 3 and 4, respectively. Among the orders detected, we found that Eurotiales represented 23%, 37%, 32% and 11% and the Mucorales: 20%, 9%, 9% and 2% of the reads in samples from Company 1, 2, 3 and 4, respectively. Other orders such as Agaricales (41%), Saccharomycetales (32%), and Wallemiales (10%) were mainly present in Company 4, 1 and 2, respectively (Figure 2b). In addition, the distribution of different fungal genera varied between companies (Figure 2c). The abundance of the genera *Kendrickiella, Sistotrema* and *Aspergillus* was significantly higher among workers in Company 3 than in other companies, while *Strobilurus, Rhodocollybia* and *Hypholoma* were mostly abundant in Company 4. OTUs with similarity to *Wallemia* genera were more common in samples from workers at Company 4 followed by 1, 2, and 3. Read abundance of genera *Wallemia, Strobilurus, Rhodocollybia* and *Hypholoma* was negatively correlated with TLR activation indicating that an increasing abundance of these genera is associated with a decreasing activation of TLRs. In contrast, genera *Candida, Rhizopus, Alternaria, Debaryomyces, Vishniacozyma* and *Mucor* were positively correlated with TLR activation (Figure S5).

Multivariate PERMANOVA and NMDS analyses (Figure 3; Table S5) revealed very similar structural patterns and showed that different recycling plants exerts strong impact on the fungal community (R^2^ = 0.44; p <0.001). Samples from workers at different companies were well separated, indicating distinct composition, and those from the same company were grouped together in ordination space (Figure 3a). We found that TLR activation variables (TLR2 and TLR4) significantly correlated with fungal community composition, with both vectors pointing in the same direction as Company 1. This indicated that species composition in samples from Company 1 induced the highest activation of TLRs, while the species composition in samples from Company 4 induced the lowest activation (Figure 3a). No correlation was found between structural patterns of fungal community and other measured vector variables (dust, endotoxin, fungal spores, actinobacteria).

**Figure 3:**
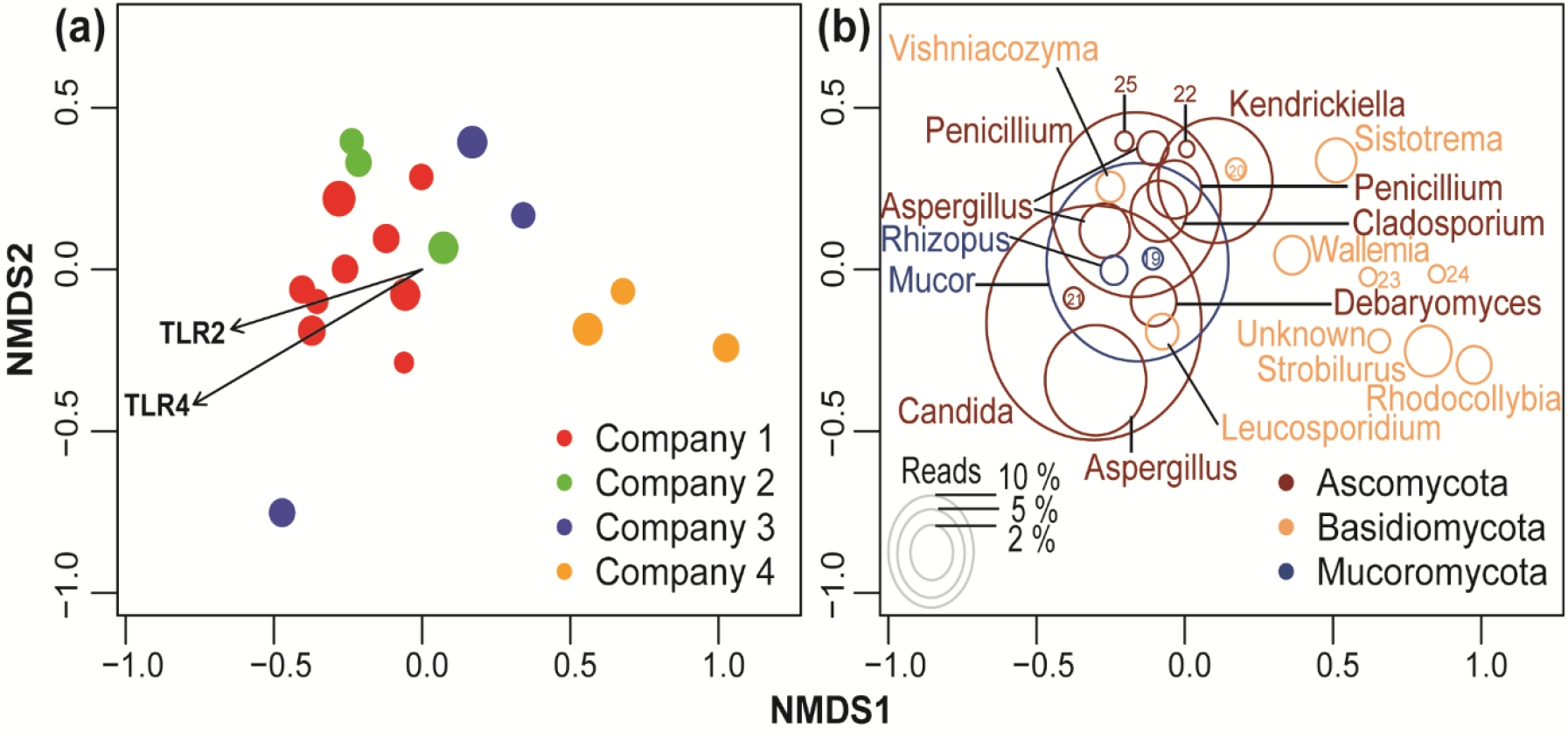
Nonmetric multidimensional scaling (NMDS) ordination analysis for fungal communities detected in personal airborne samples from workers in the four recycling plants. (a) Plot represents samples color coded by company 1-4. Vectors (TLR2 and TLR4 activation variables) shown with arrow had significant effects (p< 0.05) on the ordination configuration. (b) Plot showing fungal OTUs composition of four recycling plants. The ordination plot is based on all fungal OTUs present, but only the most common 25 OTUs are shown here, which accounted for 66% of the total reads. The number codes correspond to: Rhizopus (19), Melamspora (20), Alternaria (21), Penicillium (22), Cyllindrobasidium (23), Hypholoma (24) and Pycnopezia (25).

The distribution of the most abundant fungal OTUs differed between samples from workers at different recycling plants (Figure 3b; Table S4). In the NMDS ordination space, OTUs matched to species *Candida glaebosa* (OTU 1), *Debaryomyces hansenii (OTU 10), Rhizopus arrhizus* (OTU 17) showed their optima in the samples from workers in Company 1. Whereas the species *Mucor plumbeus* (OTU 2) and *Alternaria molesta* (OTU 23) showed quite similar distribution in companies 1 and 3, OTU 3 (*Penicillium aethiopicum*), OTU 4 (*Kendrickiella phycomyces*), OTU 5 (*Aspergillus fumigatus*) and OTU 8 (*Aspergillus flavus*) showed their highest abundance in Company 3. Only OTU 16 (*Vishniacozyma victoriae*) was more common in Company 2, while OTU 6 (*Cladosporium delicatulum*) and OTU 9 (*Strobilurus esculentus*) had species optima in Company 4 samples. Indeed, *S. esculentus* and *Rhodocollybia buryracea* were found almost exclusively in Company 4 (99.1% and 99.7% of the reads, respectively), and *Sistotrema conronilla* and *Melampsora epitea* were found almost exclusively in Company 3 (99.9% and 95.5% of the reads, respectively) (Table S4). The OTU with the highest overall percentage of reads was *Candida glaebosa* (10.1% of total reads) but was most prevalent in Company 1 (87.7 % of the reads), while reads in the other companies were 6% or less. Company 3 had median to high read percentage of most of the top 20 OTUs (Table S4).

## 5. Discussion

In this study, we characterized the occupational exposure to fungi by spores and OUT composition, endotoxins and actinobacteria present in airborne dust from waste sorting plants. The dust’s potential to induce NFkB through the activation of TLR2 and TLR4 was also assessed.

The overall dust exposure measured in this study (range: 0 – 50.2 mg m^−3^) is consistent with the concentrations measured in the waste sector in South Africa (47), Poland (48), and the exposure levels reviewed by Pearson and colleagues in (0.1-56.14 mg m^−3^) (49). Since symptoms and lung function decline have been associated with even lower dust concentrations (0.1 – 2.1 mg m^−3^) in workers at Norwegian waste recycling plants (2, 50, 51), exposure levels in the present study may be of concern. The dust exposure levels were higher than those measured in Dutch waste management plants (<0.2 – 9.1 mg m^−3^) (4, 52) as well as in Great Britain (0.07 – 17.73 mg m^−3^) (53), South Korea (0.05 – 4.51 mg m^−3^) (54) and in a newly built Norwegian waste treatment plant (0.51 – 18.93 mg m^−3^) (27). Three of the 18 workers were exposed to dust levels higher than the occupational exposure limit for total organic dust in Norway of 5 mg m^−3^ (55). The relatively high dust levels could be attributed to the fact that higher volume and different types of waste materials are recycled today than in the past. Storage of the waste materials prior to processing is likely to promote the growth of sporulating bacteria and fungi, that contribute to airborne organic dust mass (56). Moreover, we suspect an increased emission of airborne particles during the loading and shredding processes which was supported by high exposure levels measured at the reception and inspection operator position (50.22 mg m^−3^), at the machine operator station (48.55 mg m^−3^) and on the machine driving post (11.04 mg m^−3^).

We found relatively high exposure to fungal spores (median: 3×10^5^ spores m^−3^; range: 0-1.5×10^6^ spores m^−3^) and which exceeds the suggested OEL of 1×10^5^ spores m^−3^ (57). These levels are comparable to exposure ranges previously reported in the waste sector (0-2×10^6^ spores m^−3^) in Norway (2) in Canada, (0.038-1.03×10^6^ spores m^−3^) (58) and in Poland (0.143-1.65×10^6^ spores m^−3^) (17). On the other hand, much lower exposure levels have been measured in samples from Danish waste transporters (about 33 times the median level) (30) and in Brazilian waste sorting plants (3.2×10^4^ spores m^−3^) (5). Higher personal exposure ranges of 0.02-110×10^6^; 93-1140×10^6^ and 0.39-5.01×10^6^ spores m^−3^ have been measured in waste treatment plants in Germany (59), in Denmark (13) and in Norway (27), respectively. Of note, we assume that 1 CFU m^−3^ equals to 10 spores m^−3^ (60) and this equivalence is an over-simplification to allow for comparisons.

Endotoxins were detected in all samples. The levels ranged from 19 to 5321 EU m^−3^ and matched previous measurements from Dutch (2 -1,900 EU m^−3^) (32, 52), British (0.8-22656 EU m^−3^) (53) and in Norwegian (223-5277EU m^−3^) (27) waste treatment plants. Relatively lower endotoxin levels were reported in waste collectors (median: 13 EU m^−3^, range: 4-183 EU m^−3^) and in composting workers (median: 3 EU m^−3^, range: 0-730 EU m^−3^) in Norway (2, 51). Conversely, higher average concentrations of endotoxins (1123 EU m^−3^; range 4-6870 EU m^−3^) were reported from municipal waste treatment facilities in South Korea (54).

Actinobacteria were also detected but only in samples from Company 1 and ranged from 4×10^3^ to 556×10^3^ actinobacteria m^−3^. These values are comparable to previous data from Norwegian waste sectors (2), but lower than the exposure levels measured in German composting plants (59).

The fungal sequence data revealed a mycobiome that included a wide spectrum of taxonomic orders belonging to the phyla Ascomycota (Eurotiales, Saccharomycetales, Helotiales, Hypocreales and Pleosporales), Basidiomycota (Wallemiales, Agaricales, Cantharellales, Capnodiales, Tremellales, Polysporales and Pucciniales) and Mucoromycota (Mucorales). This mycobiome data is similar to that characterized in French waste recycling sector (8). The 20 most abundant OTUs (Table S4) matched species belonging to the genera *Candida* (*Debaryomyces*), *Aspergillus, Penicillium, Kendrickiella, Cladosporium, Rhizopus*, Strobilurus, Debaryomyces, *Wallemia*, Rhodocollybia, *Wishniacozyma* and *Alternaria*. We identified opportunistic pathogens and mycotoxin producers like *A. fumigatus* and *A. flavus* and allergenic species such as *Penicillium, Cladosporium, Alternaria* and *Wallemia* (61). The percentage of reads showed that *Candida (*including *C. glaebosa (*Telomorph: *Citermyces matritensis)* and *Debaryomyces hansenii (*Anamorph: *C. famata))* was the most prevalent genus (12.2%). *C. glaebosa* and *D. hansenii* are cryotolerant and halophile (salinity up to 24%) yeasts of marine origin, commonly found in sausage and dairy products (62–64). Their presence in samples from all companies indicates probable contamination of the waste materials with food waste from the dairy or sausage industry. Clinically, *C. glaebosa* and *C. famata* have been reported as new-emerging opportunistic pathogens and have been associated with invasive candidiasis (0.2-2%) in immunocompromised individuals (65, 66).

*Aspergillus* was the second most common genus and include *A. fumigatus* and *A. flavus*. These two species belong to the biosafety level 2 i.e. they can cause opportunistic infections in people with compromised immune systems (67). They are also known as causative agents of aspergillosis, and producer of aflatoxin and gliotoxin (68). The *Rhizopus* genus includes species like *Rhizopus microsporus* and *R. arrhizus* (*oryzae*) that belong to biosafety level 2 group and are classified as facultative causative agents of mucoromycosis (69). *Wallemia murae* is associated with farmer lungs and bronchial asthma in farmers (70) while *Vishniacozyma victoriae* is associated with asthma in dwellings (71).

The fungal diversity in the dust samples from the waste sorting plants varied widely between companies and to some extent reflected the potential source of waste processed during our sampling campaign. For example, *Strobilurus esculentus* is widespread on the branches and trunks of conifers, while *Rhodocollybia butyraceae*, a mushroom native to forest floors where it is common. Another example is *Melamspora epitea*, a plant leaf parasite (rust fungi). These three species were specifically characterized in samples collected at Company 4 (>95% of the reads for these species) and were indicative of the types of materials that Company 4 primarily processed during the sampling campaign.

A divergent relationship between fungal species abundance and activation of TLR2 and TLR4 was observed. Increasing abundance of *Candida, Debaryomyces, Mucor, Rhizopus, Wishnacozyma* and *Alternaria* was associated with increased activation of TLR2 and TLR4, while increasing abundance of *Penicillium, Kendrickiella, Sistotrema, Cladosporium, Wallemia, Strobilurus, Leucosporidium, Rhodocollybia, Hypholoma, Aspergillus and Melamspora* induced decreasing activation of TLR2 and TLR4 (Figure S5). Furthermore, the overall fungal community composition, measured as OTU richness was negatively correlated with TLR4 activation (Figure S4) in contrast to the fungal spore counts, which were positively correlated with TLR2 and TLR4 (Table S2). To the best of our knowledge, this is the first study to describe such a divergent relationship between biological response in in-vitro cell system with fungal exposure in a waste-sorting environment. A possible explanation of the negative association could be related to the specificity and affinity of TLR4 for O-linked mannans/ glycoproteins of fungal spore wall component (72). Because the distribution and the cell wall concentrations of these components can vary between species (73), it is likely that the increasing prevalence of species with reduced concentrations or masked O-glycoproteins may contribute to a decrease in the TLR4 activation. In particular, *Candida* yeasts have glycoproteins exposed directly on the cell wall surface in contrast to *Aspergillus* spores whose glycoprotein layer is masked by melanin and hydrophobin layers. For the spore numbers, the positive correlation with TLR2 and TLR4 could be explained by the dominance of spore production by a few sporulating fungi. Thus, increasing spore levels, but only from a few species, will increase levels of phosphor-lipomannan which activates TLR2 (74) and O-linked glycoproteins which activate TLR4 (72). Spore levels did not correlate with OTU richness, Shannon index or evenness (data not shown) but showed a negative trend relationship comparable to data reported on fungal diversity in the sawmills (75). We can also draw a parallel between these findings and the results from Viegas and colleagues who found negative trend relationships between inflammatory cytokine levels and bacterial and fungal diversity in dust collected in the Portuguese waste recycling sector (9), considering that the production of inflammatory cytokines is to some extent intrinsically linked to NFkB induction through TLRs activation. The underlaying mechanisms of this negative relationship remains unclear, however we can speculate on the pro-inflammatory properties of large insoluble beta-glucan components versus the anti-inflammatory effects of soluble beta glycans (76). As such, increasing levels of soluble beta-glycans is likely linked to increasing abundance for fungal species.

Compared to previously reported levels of bioaerosols in waste management, our results indicate moderate to high exposure levels of dust, endotoxins, and fungal spores, all of which moderately to weakly correlate with NFkB induced SEAP production through TLR2 and TLR4 activation. We found a strong correlation between total dust levels and TLR2 and TLR4 activation suggesting that dust particles carry many ligands and/or agonists for these receptors. Although both TLR2 and TLR4 were activated by the dust, the NFkB-induced response was highest with TLR2 and in most samples (16 out of 18) (Figure S2). This indicates that the dust contained more TLR2 ligands to which chronic exposure may shape the adaptive immune response toward Th2 and Treg cell differentiation pathways characterized by allergic immune responses (77, 78). Activation of TLR4 indicates the presence of ligands capable of promoting an adaptive immune response towards Th1 and Th17 differentiation, especially under chronic exposure related activation (79).

The endotoxin exposure levels were mainly in the range of concentrations that can cause inflammation (100 EU m^−3^), systemic effects (1000 EU m^−3^) or toxic pneumonitis (2000 EU m^−3^) (53). Even relatively low exposure levels in waste workers have been shown to be positively associated with inflammatory markers such as myeloperoxidase (2, 51). The correlation between TLR4 activation and the endotoxin levels was surprisingly weak albeit significant (R^2^ adjusted 0.20; p=0.03). The presence of different types of endotoxins with different levels of acylation of lipid A may explain this. Low acylated lipid A forms of endotoxins, in contrast to hexa-acylated forms, are poor stimulators of the TLR4/MD-2 complex (80).

The initial activation of PRRs by PAMPs present in the dust, followed by induction of immunological responses play a fundamental role in the development of adaptive inflammatory responses. Chronic exposure with repetitive oxidative and inflammatory reactions is associated with chronic diseases and a decline in lung function over time (18). By characterizing the potential of inhalable dust to induce NFkB responses through TLRs activation as immunological key events that potentially lead to adverse health outcomes, the present study provides a useful biological endpoint approach to consider in risk assessment and future epidemiological studies.

The significant variation in fungal spore levels and fungal species diversity between waste sorting plants suggests that different exposure-related health effects can be expected among waste company employees. Therefore, fungal exposure assessments, both by the company as well as by tasks can be crucial for evaluating health risks. Compared to the extended characterization of suspected dust components, this is an improvement in exposure and hazard characterization towards parameters that are mechanistically closer to potential health effects. Further epidemiological studies linking key immunological events, as reported here, and quantified biomarkers of adverse health outcomes in workers are needed to confirm such an association. Despite the new insights into exposure and hazard characterization from in vitro activation of TLR2 and TLR4, the small sample number in this study is a limitation. Further studies with a larger exposure sample size using TLR activation by dust from the waste management sector are needed for more definitive conclusions.

## 6. Conclusions

Besides the exposure characterization to dust, fungal spores and OTU diversity, endotoxins and actinobacteria in Norwegian waste recycling plants, the present study provided new insights into the immunological key mechanisms related to fungal exposure. Overall, workers were exposed to potentially hazardous levels of dust, endotoxins and fungal spores. A multifactor-associated NFkB-induced response through the activation of TLR2 and TLR4 by the collected dust provided a measure of the potential immunological risk effect by considering all potential hazardous contents including the fungal spore levels and diversity as well as endotoxins, bacteria and other uncharacterized components. The positive correlation of NFkB-induced cell signaling to fungal spore levels in contrast to the negative correlation to species diversity suggests that fungal spore levels as such contribute only partially to the key immunological outcomes and that identification at species level is crucial to assess exposure related health effects.

Future hazard characterization studies in waste management and other similar environments need to focus on the role of the complex composition of dust and associated health effects on workers so as to gain a more holistic view of the exposure-response relationships leading to disease.

## 7. Acknowledgments

This work was financially supported by the Norwegian Confederation of Industry (Norsk Industri) and the National Institute of Occupational Health (STAMI). We thank dr. Kari Kulvik Heldal for advice and support.

